# The distance between the plasma membrane and the actomyosin cortex acts as a nanogate to control cell surface mechanics

**DOI:** 10.1101/2023.01.31.526409

**Authors:** Sergio Lembo, Léanne Strauss, Dorothy Cheng, Joseph Vermeil, Marc Siggel, Mauricio Toro-Nahuelpan, Chii Jou Chan, Jan Kosinski, Matthieu Piel, Olivia Du Roure, Julien Heuvingh, Julia Mahamid, Alba Diz-Muñoz

**Author notes:** Current address: Institute of Science and Technology Austria (ISTA), Klosterneuburg, Austria. Current address: Mechanobiology Institute, Department of Biological Sciences, National University of Singapore, Singapore (Chii Jou Chan). These authors contributed equally to this work (Sergio Lembo, Léanne Strauss). Collaboration for joint PhD degree between EMBL and Heidelberg University, Faculty of Biosciences.

## Abstract

Animal cell shape changes are controlled by the actomyosin cortex, a peripheral actin network tethered to the plasma membrane by membrane-to-cortex attachment (MCA) proteins. Previous studies have focused on how myosin motors or actin turnover can generate the local deformations required for morphogenesis. However, how the cell controls local actin nucleation remains poorly understood. By combining molecular engineering with biophysical approaches and *in situ* characterization of cortical actin network architecture, we show that membrane-to-cortex tethering determines the distance between the plasma membrane and the actomyosin cortex at the nanoscale of single actin nucleators. In turn, the size of this gap dictates actin filament production and the mechanical properties of the cell surface. Specifically, it tunes formin activity, controlling actin bundling and cortical tension. Our study defines the membrane-to-cortex distance as a nanogate that cells can open or close by MCA proteins to control the activity of key molecules at the cell surface.

---

The cell cortex is a thin network of actin filaments, myosin motors and actin-binding proteins linked to the plasma membrane^1^. Spatial regulation of cortex composition and mechanics drives cell shape changes during key physiological processes such as division, migration and differentiation. Thus, deciphering how cells control cortex mechanics locally is essential to understand the remarkable ability of animal cells to dynamically change their shape. To identify the molecular mechanisms which spatially control cortex mechanics, studies have focused on the distribution of active myosin molecules^2,3^. However, how the cell regulates local actin nucleation activity, which determines the local architecture of the cortical network, remains poorly understood.

The cell cortex is tethered to the plasma membrane by membrane-to-cortex attachment (MCA) proteins. Canonical MCA proteins, such as the ERM family (ezrin, radixin and moesin), are key regulators of membrane mechanics, in particular membrane tension^4^. Activated by phosphorylation, ERM proteins distribute asymmetrically during many morphogenetic processes, coinciding with mechanically and functionally distinct cortical regions in cells. For example, they specifically localize and regulate the rear of migrating cells^5^, the cytokinetic ring of dividing cells^6,7^, the microvillar domain of mouse oocytes^8^, and the apical domain of early mouse blastomeres^9,10^ and epithelial cells^11,12^. Thus, we wondered whether and how membrane-to-cortex tethering controls cortical mechanics.

In this study, we find that reducing the distance between the plasma membrane and the most peripheral actin filaments leads to a reduction in cortical tension and thickness. By employing *in situ* cryo-electron tomography, we reveal changes at the level of single actin filaments in the architecture of the cortical network. Finally, by using chemical perturbations and engineered MCA proteins of different lengths, we find that membrane-to-cortex distance controls actin nucleation by formins. Together, these results identify MCA proteins as major regulator of the physical space between the plasma membrane and the actin cortex, which in turn controls key molecular activities at the cell surface.

## Membrane-to-cortex tethering regulates cortical tension and thickness

ERM proteins are enriched at functionally distinct regions of the cell cortex with differential mechanics. Thus, we examined whether membrane-to-cortex tethering has a direct effect on the mechanical properties of the cortex. To keep a constant cell geometry between our measurements, and induce the formation of a uniform cortical network^13^, we plated cells on circular micropatterns, such that they uniformly adapt into a dome-like shape (**Fig. 1a**). We then increased membrane-to-cortex tethering first by expressing a constitutively active version of ezrin (CAezrin, T567D^14^) in a doxycycline inducible manner in NIH 3T3 fibroblasts (**Fig. 1b** top and **Extended Data Fig. 1a,b**). Cortical tension, a key mechanical property of the cortex, was measured by atomic force nano-indentation (**Fig. 1a, Extended Data Fig. 1c**). Increasing membrane-to-cortex tethering by CAezrin expression led to a decrease in cortical tension (**Fig. 1b** bottom). To rule out an effect of ezrin’s signaling rather than its tethering function (reviewed in^15^), we next used our recently developed iMC-linker, which is a synthetic molecular tool that mimics the domain architecture of endogenous MCA proteins but is inert regarding signaling^16^. Specifically, the iMC-linker is reduced to three small protein domains: the CH1/CH2 domain of Utrophin^17^ to bind cortical actin, the lipidation motif of Lyn^18^ to insert into the plasma membrane, and a fluorescent protein to link the two domains and visualize its subcellular localization (**Extended Data Fig. 1d**). Consistent with the effect of CAezrin expression, increasing tethering by expression of the iMC-linker (**Extended Data Fig. 1e**) also significantly decreased cortical tension in fibroblasts in a density-dependent manner (**Fig. 1c** bottom, **d**). Thus, MCA proteins not only connect the plasma membrane and the cortex, but also regulate cortical tension. To rule out a change in actin dynamics by the actin-binding activity of the iMC-linker, we expressed the actin-binding domain alone and found that it was not sufficient to decrease cortical tension (**Extended Data Fig. 1f**), demonstrating that the tethering of cortical actin to the membrane is key for this mechanical phenotype. Next, to rule out any effects due to potential differences in cell adhesion^19^, we measured cortical tension by micropipette aspiration of cells in suspension. Regardless of cell adhesion to the substrate, iMC-linker expressing cells had lower cortical tension (**Extended Data Fig. 1g**). Finally, we overexpressed CAezrin or the iMC-linker in naïve mouse embryonic stem cells (mESCs). Nanoindentation measurements consistently showed a decrease in cortical tension (**Fig. 1 e,f, Extended Data Fig. 1h-j**), which demonstrates that regulation of cortical tension by membrane-to-cortex tethering is a consistent feature among different cell types.

**Fig. 1.**
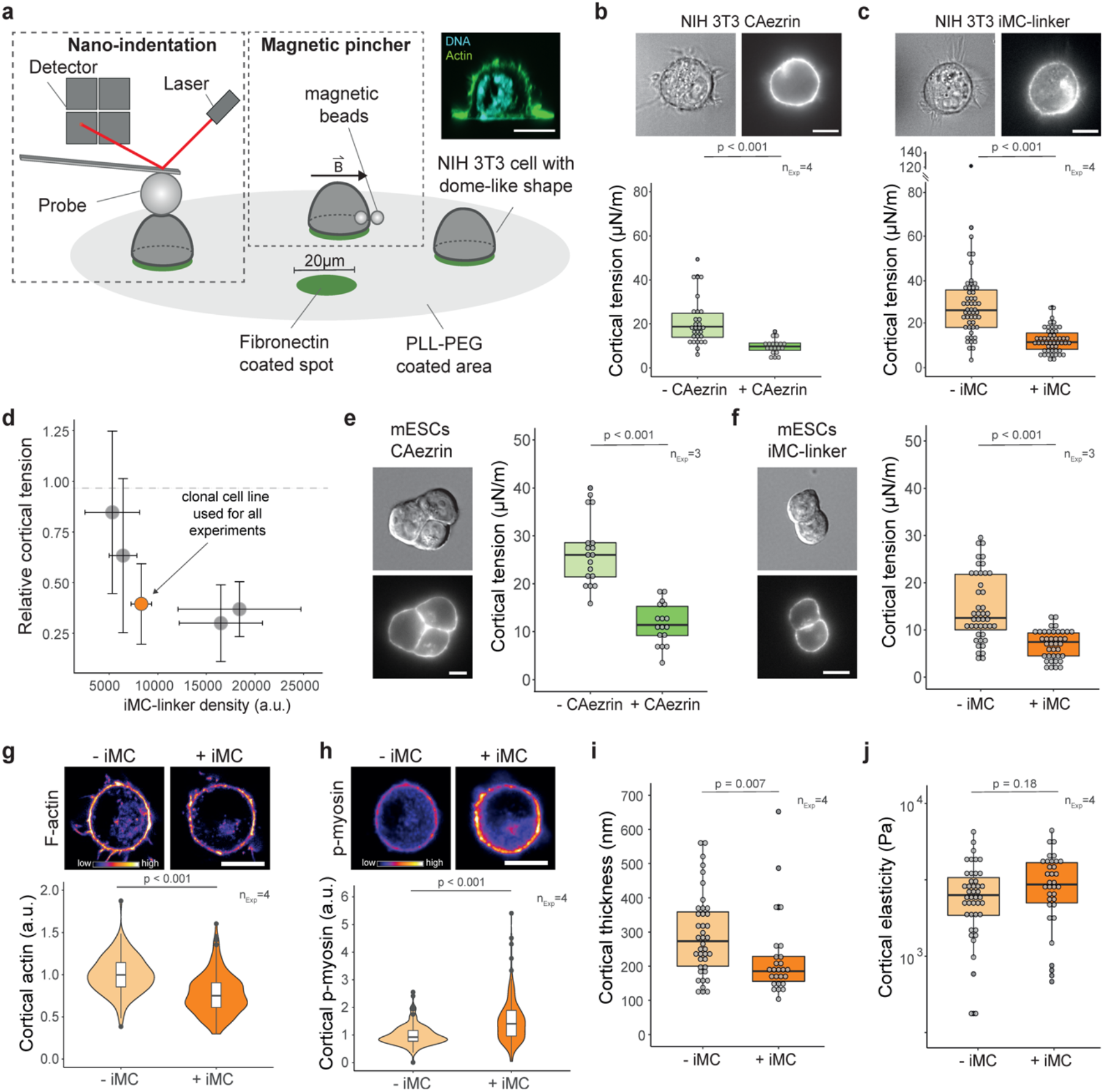
Membrane-to-cortex tethering regulates cortical tension and thickness. **a**, Schematic representation of the experimental setup for cortical tension and thickness measurements of NIH 3T3 fibroblasts, by nano-indentation and a magnetic pincher (vector B depicts the magnetic field), respectively. Cells are seeded on micropatterned dishes to acquire pseudo-spherical shape. Insert: representative xz view of a cell fixed and stained with phalloidin (F-actin) and DAPI (DNA). **b,c**, (top) Representative bright field and epifluorescence images of NIH 3T3 fibroblasts expressing CAezrin (b) or iMC-linker (c). (bottom) Cortical tension of NIH 3T3 fibroblasts upon expression of the respective construct. **d**, Relative cortical tension of various iMC-linker expressing clonal NIH 3T3 fibroblasts lines. Values are normalized to the respective non-induced control. Orange highlights the clonal line used in the rest of this study, others are depicted in grey. Each dot represents the median of independent experiments of a single clonal line (n_Exp_ ≥ 3). **e,f**, (left) Representative differential interference contrast and epifluorescence images of a mESCs expressing CAezrin (e) or iMC-linker (f). (right) Cortical tension of NIH 3T3 fibroblasts upon expression of the respective construct. **g,h**, (top) Representative confocal images of NIH 3T3 fibroblasts seeded on micropatterned dishes, and stained with phalloidin (F-actin) (g) and an anti-p-myosin antibody (h). (bottom) Normalized mean fluorescence intensity of cortical F-actin (g) and p-myosin (h). **i**,**j**, Cortical thickness (i) and cortical elasticity (j) in NIH 3T3 fibroblasts upon expression of the iMC-linker. Each dot represents the mean of multiple measurements of a single cell unless specified otherwise. Scale bars = 10 μm. n_Exp_ = number of independent experiments, a.u. = arbitrary units. Normality of data distribution was tested by Shapiro-Wilk test. Two-tailed t-test was used for normally distributed data. Otherwise, a non-parametric Wilcox test was used.

A decrease in tension could result from reduced myosin-2 activity or a change in the architecture of the cortical actin network^3,20^. First, we quantified the relative concentration of phosphorylated myosin-2 (p-myosin) at the cell surface. Surprisingly, we found a clear increase in the amount of p-myosin at the cell periphery (**Fig. 1h, Extended Data Fig. 1k** right), which would instead be expected to increase tension. Next, as a first-order readout of cortical architecture, we measured cortical thickness and elasticity using a magnetic pincher^21^. Cortical thickness was reduced by 30% upon iMC-linker expression (**Fig. 1i**), without a significant change in elasticity (**Fig. 1j**). Moreover, we observed a slight decrease in the relative amounts of filamentous actin (F-actin) at the cortex (**Fig. 1g, Extended Data Fig. 1k** left). Together, our results suggest that increasing membrane-to-cortex tethering reduces cortical tension by changing cortical network architecture.

## Membrane-to-cortex tethering regulates cortical network architecture and the membrane-to-cortex distance

Analyzing actin network architecture in cells is a long-standing challenge because actin filaments are poorly preserved during chemical fixation and in traditional room temperature microscopy methods^22^. Moreover, the filaments are 7 nm thick and thus require high resolution imaging. The combination of light and electron microscopy approaches have shed some light on the architecture of the actin cytoskeleton at the basal surface and migration front in some cell types^23,24–27^. Yet, such methods have failed to provide a characterization of the cell cortex, which is further hampered by its diffraction-limited thickness^20,28^, close proximity to the membrane, and high density of actin filaments^29^.

To assess how membrane-to-cortex tethering affects the architecture of the cortical actin network, we established a cryo-electron tomography (cryo-ET) workflow, which can resolve individual actin filaments *in situ*. In alignment with our measurements of cell surface mechanics above, NIH 3T3 fibroblasts were seeded on micropatterned grids. To target the cortex at the rounded parts of the cell, we used focused ion beam milling^30^ (FIB) to produce ~200 nm thin lamellas in the equatorial plane. To ensure structural preservation of this area and stability of the lamellae, the lamellae were generated across two vitrified cells growing on adjacent micropatterns^31^ without forming cell-cell adhesions (**Extended Data Fig. 2a-e**). We obtained 3D cryo-ET reconstructions at 1.3 nm/pixel, semi-automatically segmented individual actin filaments and the plasma membrane, and extracted structural parameters of the actin network (**Fig. 2a-d, Extended Data Fig. 2f, Extended Data Fig. 3a, Supplementary Videos 1-4**): our ultrastructural analysis showed that increased membrane-to-cortex tethering by expression of the iMC-linker leads to a reduction in the total amount of actin in filaments (**Fig. 2b**), in agreement with fluorescence-based quantification of cortical F-actin amount (**Fig. 1g**) and thickness (**Fig. 1i**). This reduction is observed especially at a distance beyond ~100 nm from the plasma membrane (**Fig. 2b**). However, the expression of the iMC-linker led to more actin filaments very close (0 to 40 nm) to the membrane compared to the control (**Fig. 2c**).

**Fig. 2.**
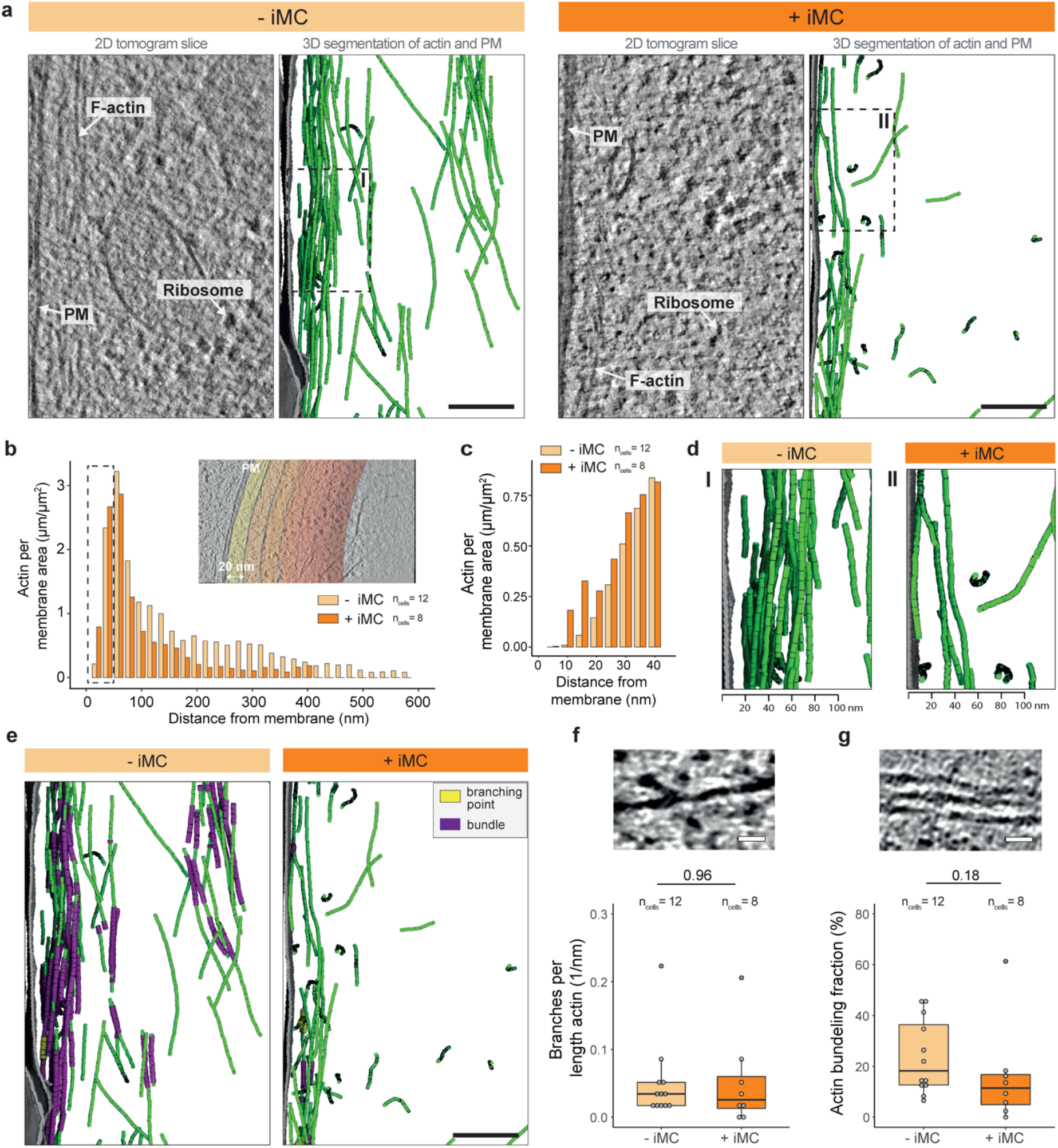
Membrane-to-cortex tethering regulates cortical architecture. **a**, Representative tomogram slices and segmentations of actin filaments (green) and plasma membrane (grey) rotated such that the membranes are aligned to the y-axis in control (left) and iMC-linker expressing (right) NIH 3T3 fibroblasts. See Supplementary Videos 1-4 for complete tomograms. Dashed squares (I) and (II) indicate the areas depicted in (d). **b**, Areal density of F-actin as a function of distance from the plasma membrane.Insert depicts representative tomogram slice with the histogram bins. **c**, Close up on the first 40 nm with 5 nm bins. **d**, Magnification from (a) showing membrane-to-cortex distance. **e**, Representative segmented tomograms shown in (a) displaying the detected actin bundles and branches in control (left) and iMC-linker expressing (right) NIH 3T3 fibroblasts (each branching point is illustrated by a segment for better visualization). **f,g**, (top) Representative image of a F-actin branching point (f) or bundle (g). (bottom) Number of F-actin branching points per actin length (f) and normalized amount of F-actin in bundles (g). Scale bars = 100 nm in (a) and (e), = 20 nm in (f) and (g). PM denotes the plasma membrane. n_cells_ = number of cells. Normality of data distribution was tested by Shapiro-Wilk test. Two-tailed t-test was used for normally distributed data. Otherwise, a non-parametric Wilcox test was used.

Next, we aimed to examine the topology of actin filaments within the cortex. Actin filaments can be organized into bundles, whose filaments are elongated by formins, or branched networks, generated by the Arp2/3 complex from preexisting filaments^32^; *in vitro*, these nucleators generate distinct network topologies with different mechanical properties^33–36^. In cells, the cortex appears soft if predominantly nucleated by the Arp2/3 complex, and bundled and stiff if predominantly generated by formins^37,38^. The activity ratio of the two nucleators can thus control the network architecture and thereby mechanics of the cell cortex^37,38^. In order to analyze whether increasing membrane-to-cortex tethering affects the topology within the cortex, we derived inter-filament orientations and distances. Actin branching was low in both control and upon expression of the iMC-linker, and was not significantly different between the two conditions (**Fig. 2e,f, Extended Data Fig. 3a,c**). We however observed a reduction in the fraction of bundled actin (**Fig. 2e,g, Extended Data Fig. 3a,b,d**).

Together, our findings allow us to understand how increased membrane-to-cortex tethering changes cortical network architecture, pointing to a specific effect on bundling, and link it to the resulting cell surface mechanics.

## Membrane-to-cortex tethering regulates the actin nucleator landscape

Our findings reveal that increasing membrane-to-cortex tethering reduces cortical thickness (**Fig. 1i**), amount of actin filaments (**Fig. 1g** and **2b**), their bundling (**Fig. 2e,g**), and the distance between the membrane and the most peripheral actin filaments (**Fig. 2c,d**). This led us to investigate actin filament polymerization at the cell surface, as our data suggests that increasing tethering might limit formin activity at the cell cortex.

mDia1 has been shown to be the primary formin at the cortex^39^. Of note, its activity is mechanosensitive and dependent on the environment^40,41^, being more processive when bound to the plasma membrane and also, upon pulling force on the polymerizing filament. First, we quantified mDia1 amount, which was unchanged by the expression of the iMC-linker (**Fig. 3a, Extended Data Fig. 4a**). Next, we tested if the effects of increased tethering on cortical mechanics are the result of reduced formin activity. To that end, we used SMIFH2, a pan-formin inhibitor ^42^. In agreement with previous reports, cortical tension was strongly reduced upon inhibitor treatment^3^ in our control cells, to strikingly similar levels as expression of the iMC-linker alone (**Fig. 3b**). Moreover, in cells with increased tethering, formin inhibition had little additional effect on cortical tension. Importantly, to rule out off-target effects of SMIFH2, which has been shown to inhibit members of the myosin superfamily^43^, we also tested the effect of direct myosin-2 inhibition. We found that Blebbistatin treatment decreases cortical tension also upon expression of the iMC-linker (**Extended Data Fig. 4b**), suggesting that our SMIFH2 findings are specific to formins. In contrast, inhibition of the Arp2/3 complex by the small molecule CK-666^44^ increased cortical tension (**Extended Data Fig. 4c**), regardless of expression of the iMC-linker. Finally, simultaneous inhibition of both cortical nucleators led to a strong reduction in cortical tension, irrespective of the level of tethering (**Extended Data Fig. 4d**).

**Fig. 3.**
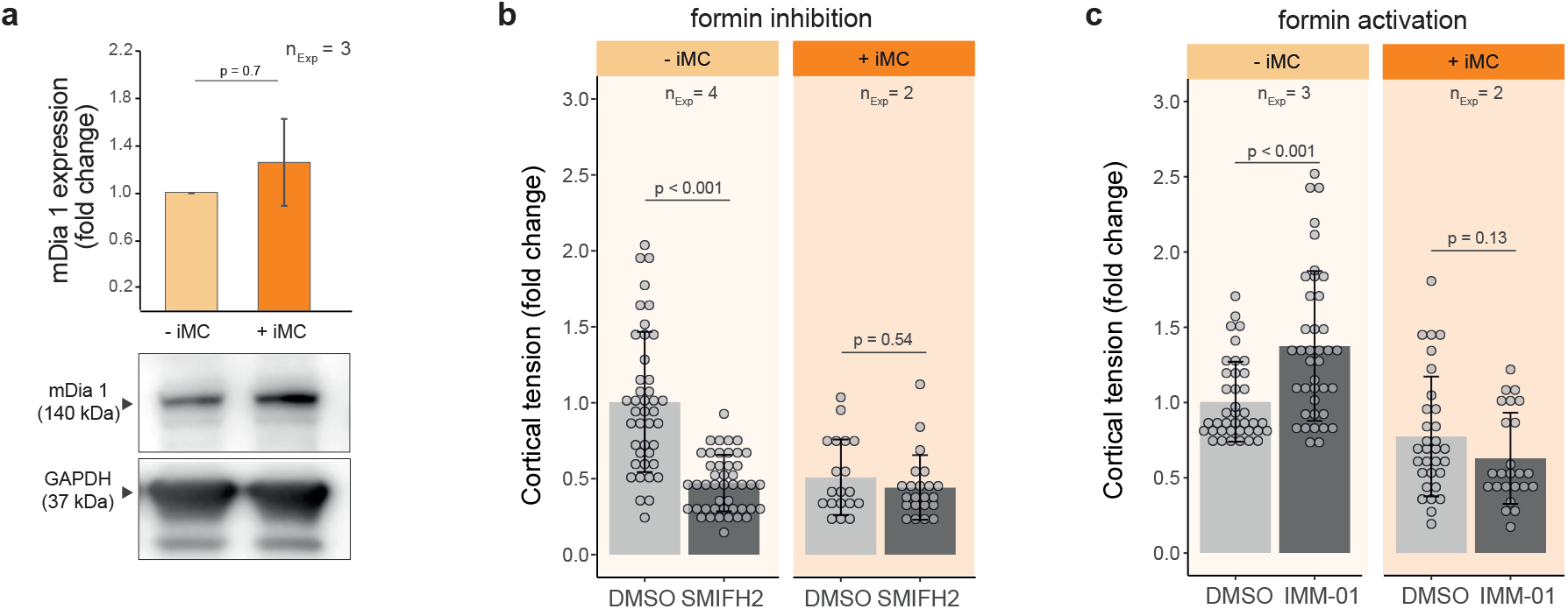
Membrane-to-cortex tethering modulates nucleation of cortical actin filaments by formins. **a**, (top) Relative formin mDia1 protein amount quantified by western blotting. Protein levels are normalized to loading control GAPDH and to protein expression of control cells. (bottom) Representative western blot bands for mDia1 and GAPDH. **b,c**, Relative cortical tension of NIH 3T3 fibroblasts (± iMC) upon treatment with SMIFH2, a pan-formin inhibitor (b) or IMM-01, a formin’s activator (c). Tension values are normalized to the mean cortical tension of control cells (-iMC) treated with vehicle only (DMSO). Each dot represents the mean of multiple measurements of a single cell. n_Exp_ = number of independent experiments. Normality of data distribution was tested by Shapiro-Wilk test. Two-tailed t-test was used for normally distributed data. Otherwise, a non-parametric Wilcox test was used.

Taken together, our findings thus far suggest that formin activity is minimal upon expression of the iMC-linker. To test if the reduction in cortical tension by increased tethering can be restored by activating formins, we used the small molecule IMM-01^45^, which locks formins in their active conformation. Such treatment increased cortical tension in control cells, but failed to do so in cells expressing the iMC-linker (**Fig. 3c**). This strongly suggests that reducing the distance between the plasma membrane and the cortex (**Fig. 2c**) precludes formin activity at the interface of these two structures.

## The membrane-to-cortex distance limits the activity of formins at the cell surface

We hypothesized that the reduction in the membrane-to-cortex distance at the nanoscale is the mechanism behind the inhibition of formins. Such “nanogating” would be expected to operate at the scale of 10-20 nm (^29^ and **Fig. 2c**). This hypothesis is based on recent cryo-electron microscopy structures indicating that formins undergo a conformational extension from 10 to 20 nm upon activation^46^. Active formin molecules would thus require larger gaps at the membrane-to-cortex interface than the more compact nucleation complex Arp2/3^47^.

To test our nanogating hypothesis, we introduced a linker with a predicted length of over 30 nm (iMC-6FP-linker), thus longer than active ezrin (25 nm^48^) and the 10 nm long original iMC-linker (**Fig. 4a**,). Although, the iMC-6FP-linker increased tethering between the membrane and the cortex (**Extended Data Fig. 5a**) and was expressed at higher levels than the original iMC-linker construct (**Extended Data Fig. 5b**), it did not lead to a reduction in cortical tension, in stark difference to the CAezrin and the original iMC-linker (**Fig. 1b,c**, and **4b**). Taken together, we propose that the membrane-to-cortex distance acts as a nanogate, that cells can locally modulate through MCA proteins, to regulate formin activity and thus cortex architecture and its mechanics (**Fig. 4c**).

**Fig. 4.**
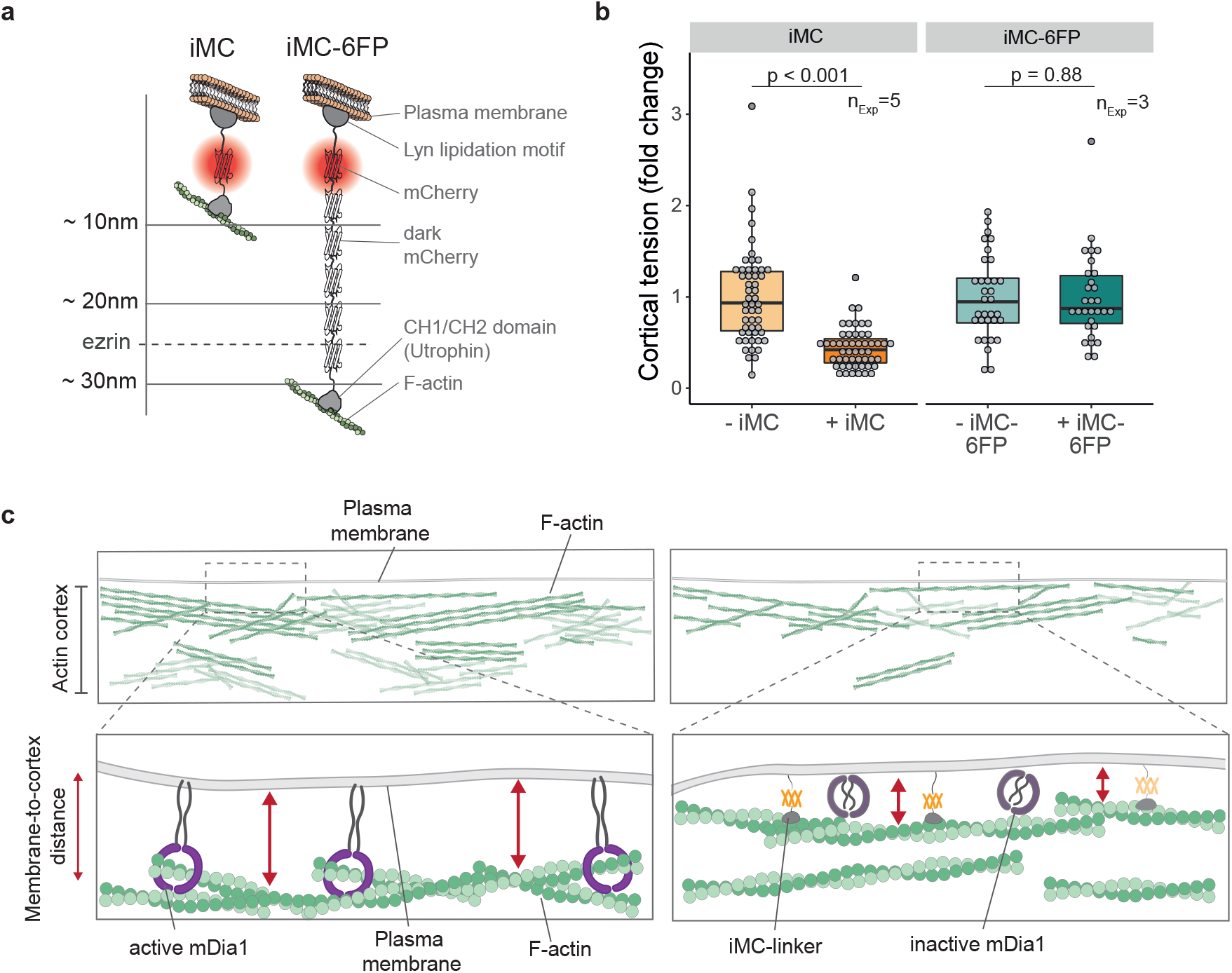
Membrane-to-cortex distance gates the activity of cortical actin regulators. **a**, Schematic representation of iMC-linker (left) and iMC-6FP-linker (right). iMC-6FP-linker construct is built with 5 dark (*i.e*. non-fluorescent) mCherry molecules in addition to domains present in the original iMC-linker. Dashed line: active ezrin length^48^. **b**, Relative cortical tension in NIH 3T3 fibroblasts upon expression of iMC-linker (normalized data from fig. 1c) or the iMC-6FP-linker, normalized to the respective control. Each dot represents the mean of multiple measurements of a single cell. n_Exp_ = number of independent experiments. **c**, Working model: membrane-to-cortex tethering regulates the membrane-to-cortex distance in a MCA protein length-dependent fashion. The iMC-linker and the endogenous MCA proteins reduce the membrane-to-cortex distance and consequently hampers the ability of mDia1 to elongate actin filaments from the plasma membrane. Normality of data distribution was tested by Shapiro-Wilk test. Two-tailed t-test was used for normally distributed data. Otherwise, a non-parametric Wilcox test was used.

## Discussion

In recent years, membrane-to-cortex tethering has gained attention, owing to its regulation of fundamental cellular processes such as stem cell differentiation^16,49^, cell division^7^ and migration^50,51^. However, how it can regulate such a diversity of cellular states remains poorly understood.

Interestingly, the spatial localization of MCA proteins often mirrors local differences in cortex mechanics, as seen for example in the apical domain of epithelial cells and early mouse blastomeres, the back of migrating cells, or the cytokinetic ring^5–10,12,52–54^. This raises the intriguing idea that a functional relation exists between these two phenomena. Here, we show that MCA proteins control cortical mechanics in a length- and density-dependent manner, limiting formin activity at the cell cortex and thus controlling cortex architecture. The redundancy of MCA proteins found in metazoan^55^ would render such a mechanism robust to genetic instability and enable control by different regulatory pathways. Moreover, this novel mechanism suggests that cells could employ gradients in linker density to regulate cortical tension, resulting in local contractions that drive cell deformations.

Membrane-to-cortex tethering limits lateral diffusion of proteins at the plasma membrane (picket fence model^56^). In this study, we show that it also regulates protein activity in the third dimension. Specifically, it regulates cortical tension by modulating formin activity, but only when expressing linkers with a predicted length shorter or equal to that of active ezrin. Together, our findings highlight how the membrane-to-cortex distance is a novel geometrical parameter that can regulate protein activity at the cell periphery. We observe the same mechanical phenotype in iMC-linker and CAezrin expressing cells, which supports an unspecific allosteric inhibition of a geometrical nature. The fact that IMM-01 is not able to increase cortical tension in cells expressing the iMC-linker further supports that the membrane-to-cortex distance by itself regulates mDia1 activity directly. Given the mechanosensitivity of formins^57–60^, and specifically that of the key formin at the cell cortex mDia1^39,58,60,61^, it is tempting to speculate that a feed-forward loop might be at play: membrane-to-cortex tethering may alter formins’ ability to efficiently elongate filament ends for significant durations^61^ at the cell surface. This is expected to lead to a reduction in cortical bundles and tension, which in turn further decreases formin-dependent filament elongation, as formins are more processive upon tension on the polymerized filament^39,41^.

Connecting cell-scale mechanics to defined cytoskeletal topologies is a current challenge in mechanobiology. There is an urgent need to develop methods that allow descriptions of cytoskeletal network architecture in quantitative terms at subcellular resolution, to guide reconstitution and modelling efforts, and interpret biophysical measurements. Although other actin structures have been resolved by cryo-ET^62–65^, we provide the first characterization of cortical architecture with single filament resolution away from the adhesive substrate in its native state. By keeping a constant cell geometry between our biophysical, biochemical and structural approaches we are able to link cell-scale mechanics to its molecular constituents.

In summary, our work identifies the membrane-to-cortex distance as a nanogate, that acts as a new and likely universal type of control switch for cortical mechanics. By deploying MCA proteins, cells can control this key geometrical parameter to pattern protein activity at the cell surface, and thereby regulate a plethora of biological processes.

## Material and methods

### Cell culture

NIH 3T3 fibroblasts (CRL-1658, ATCC repository) were cultured in Dulbecco’s Modified Eagle Medium (DMEM) with 4.5 g/l D-glucose (Merck), supplemented with 10% fetal bovine serum and 1% Penicillin-Streptomycin (10,000 U/ml). Mouse Embryonic Stem Cells (mESCs, kindly provided by the Austin Smith’s group, Cambridge Stem Cell Institute), were maintained in serum-free N2B27 medium containing 2i (1 mM PD0325901and 3 mM CHIR99021, both Tocris) and LIF (10 mg/ml, Merck) on polystyrene culture dishes coated with 0.1% (w/v) Gelatin (Merck) solution. N2B27 medium was prepared from a 1:1 mixture of DMEM/F12 (without HEPES, with L-glutamine) and neurobasal medium (no L-glutamine), supplemented with 0.5x B-27 (without vitamin A) and 0.5x N-2 supplement, 2.5 mM L-glutamine, 10 mg/ml BSA fraction,10 mg/ml human recombinant insulin and 1% Penicillin-Streptomycin (10,000 U/ml). Cells were cultured at 37° C with 5% CO_2_ on polystyrene 100 mm dishes (Corning). Cells were passaged every 2-3 days (up to 70% confluency) using 0.05% Trypsin-EDTA. Cells were routinely tested for mycoplasma contamination using MycoAlert detection kit (Lonza) according to manufacturer’s instruction.

All the transgenic cell lines used in this study (**Supplementary Table 1**) were generated using a PiggyBac transposon system^66^. The gene of interest was cloned via Gibson assembly in a custom PiggyBac vector (kindly provided by Jamie Hackett’s group, EMBL Rome) downstream of a Doxycycline (dox)-responsive TRE3G promoter. The transposable region of the vector contained also a TET3G and a Neomycin resistance under the control of a CAG promoter. Stable cell lines were obtained by co-transfecting the PiggyBac vector and the PiggyBac transposase using lipofectamine 3000. Successfully transfected cells were then positively selected by adding 1.2 mg/ml Geneticin to the medium. Expression of the constructs was induced by the addition of 1 μg/ml dox to the medium 16 hrs before probing the cells.

Monoclonal cell lines were finally generated by limiting dilution, and screened via flow cytometry for low background expression and similar expression upon dox-dependent induction (**Extended Data Fig. 5a**).

All the reagents were purchased from Thermo Fisher Scientific unless overwise specified.

### iMC-6FP-linker cloning

iMC-6FP-linker was generated by introducing five non-fluorescent mCherry proteins (dark mCherry, Y72S) between the fluorescent mCherry molecule and the actin binding domain of the iMC-linker ^67^. Single dark mCherry molecules were added using Type IIS endonucleases cloning ^68^. The length of the molecule was predicted based on the crystal structure of a single mCherry molecule.

### Cellular micropatterning on glass

NIH 3T3 fibroblasts were seeded on 20 μm adhesive spots to unify their geometry and ensure a spherical shape. For micropatterning of the dishes, an inverted Ti2^®^ Nikon microscope equipped with the Primo micropatterning module (Alvéole) was used. In brief, 35 mm glass-bottomed dishes were plasma treated for 1 min and a small PDMS stencil with an inner hole of ~10-20 μl was placed in the middle of the dish to reduce reagents usage. Next, the glass surface inside the stencil was coated with a non-adhesive layer of 0.2 mg/ml polyethylene glycol (PLLg-PEG, Susos AG) for 1 hour at RT. Then, the surface was washed trice with PBS before applying a thin layer of 4-benzoylbenzyl-trimethylammonium chloride micropatterning reagent (PLPP, Alvéole). Using a 375 nm laser (4.5 mW) 20 μm circles spaced by 40 μm from each other were photo-patterned onto the glass surface, using the μmanager software v.1.4.22 with the Leonardo plugin v.4.12 (Alvéole). Finally, the dish was coated with 50 μg/ml human fibronectin (Corning) for 30 min and 10^5^ cells were seeded onto the patterned dish. Cells were allowed to settle and adhere for 20 min before non-attached cells were washed out. Experiments were performed 2 hrs after cell seeding if not indicated otherwise.

### Single-cell atomic force spectroscopy

Tether extrusion and nano-indentation were performed on a CellHesion^®^ 200 atomic force spectrometer (Bruker) integrated into an Eclipse Ti^®^ inverted light microscope (Nikon). Measurements were run at 37 ̊ C with 5% CO_2_ and samples were used no longer than 1 hour for data acquisition. Measurements were acquired at a sampling rate of 2 kHz and analyzed using the JPK Data Processing Software.

<u>Apparent plasma membrane tension</u> (T_(app)_) (the sum of in-plane tension (T_m_) and the adhesion energy resulting from membrane-to-cortex tethering (γ)) depends on the breakage tether force (f_0_) and the bending rigidity of the membrane (k)^69^, which we assumed is unchanged between our experimental conditions.

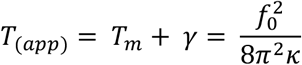

To measure the breakage tether force, we used OBL-10 Cantilevers (spring constant ~0.06 N/m; Bruker) calibrated using the thermal noise method and coated for 1 hour at 37 ̊ C with 2.5 mg/ml Concanavalin A (Thermo Fischer Scientific), which binds polysaccharides expressed on the surface of the cell. Before the measurements, cantilevers were rinsed in PBS. NIH 3T3 fibroblasts were washed and probed in Dulbecco’s Modified Eagle Medium (DMEM) with 4.5 g/l D-glucose (Merck) supplemented with 2% fetal bovine serum (Thermo Fisher Scientific) and 1% penicillin-streptomycin (10,000 U/ml)(Thermo Fisher Scientific). mESCs were probed in growth medium.

In brief, approach velocity was set to 0.5 μm/s while contact force and contact time ranged between 100 to 200 pN and 100 ms to 10 s, respectively, in order to maximize the probability to extrude single tethers, while minimizing the experimental stress on the cells. After contacting the cell surface, the cantilever was retracted for 10 μm at a velocity of 10 μm/s. The position was then kept constant for ~30 s and tether force was recorded at the moment of tether breakage. Each cell was probed for a maximum of 10 attempts or until 3 single tethers were successfully extruded. Tether force values from a single cell were then averaged.

For <u>cortical tension measurements on adherent cells</u>, nano-indentation was performed using tipless MLCT-O10 type C cantilevers (spring constant ~0.01 N/m; Bruker), calibrated using the thermal noise method. A silica bead with a diameter of 10 μm (Microparticles) was glued onto the cantilever using a 2-part epoxy resin (UHU) with 5 min working time. Upon full hardening of the glue, the cantilever was coated for 30 min at 37 ̊ C with a 1% (w/v) Pluronic F-68 (Thermo Fisher Scientific) in milli-Q water and rinsed with PBS, aiming to prevent non-specific adhesion to the probed cells. Force-distance curves were acquired using 500 pN contact force and 0.4 μm/s approach/retract velocity. Up to 5 curves, were taken per cell, with a 10 s waiting time between successive curves to prevent history effects.

Cortical tension (T_c_) was determined by fitting each force-indentation curve between 0 (the contact point) and 300 nm with a liquid droplet model suited for nano-indentation experiments^70^:

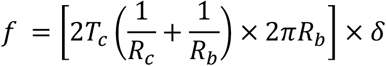

Where f is the force applied to the cell surface which leads to a displacement (δ) of cellular material; R_c_ and R_b_ are the radius of the cell and the silica bead glued on the cantilever, respectively.

### Micropipette aspiration

Micropipettes were forged by pulling glass capillaries (World Precision Instruments TW100-3) using a P-97 Flaming Brown needle puller (Sutter Instrument) in order to obtain a radius (R_p_) of 4-6 μm. The micropipette was mounted on a micromanipulator (Narishige, MON202-D) and connected to a microfluidic pump (Fluigent, MFCS) delivering negative pressures with a 7 Pa resolution. The pressure is controlled using custom-made Labview (National Instruments) software.

To measure cortical tension, NIH 3T3 fibroblasts were detached from the substrate using 0.05% Trypsin-EDTA (Thermo Fisher Scientific), seeded on a 5 cm glass-bottom dish pre-coated with a non-adhesive layer of 0.2 mg/ml polyethylene glycol (PLLg-PEG, Susos AG) and incubated at 37° C with 5% CO_2_ for 4 hrs to allow cell shape recovery. After the incubation, cells were probed by bringing the micropipette in contact, and increasing stepwise a negative pressure until a deformation of the cells with the same radius of the micropipette (R_p_) was formed. At steady state, the surface tension (T_c_) of the blastomeres was calculated based on Young–Laplace’s law^71^:

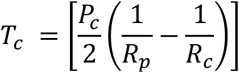

Where P_c_ is the pressure used to deform the cell of radius (R_c_).

### Drug treatments

NIH 3T3 fibroblasts were seeded on micropatterned dishes as described above and actomyosin perturbing drugs were added directly to the medium to reach a final concentration of 40 μM for SMIFH2, 100 μM for CK-666, 25 μM for IMM-01 and 20 μM for (−)-Blebbistatin (all Merck). Cells were then incubated with the drug for 1 hour before probing with an atomic force spectrometer in presence of the drug. Control cells were treated with vehicle only (DMSO) at the same final concentration of the respective drug.

### Magnetic pincher

24-48 hrs before magnetic pinching experiments, cells were seeded at a density of 5*10^3^ cells/cm^2^ in the presence of superparamagnetic beads (M-450, Dynabeads) at a density of 10^4^ beads/cm^2^.

On the day of the experiment, cells were detached from the substrate using 0.05% Trypsin-EDTA (Thermo Fisher Scientific) and plated on micropatterned dishes as described above. Then 1.5*10^5^ M450 beads were added to the dish, and the medium was supplemented by 20 mM of HEPES buffer, to mitigate pH fluctuations during the experiment.

The magnetic pincher setup is mounted on an Axio A1 inverted microscope (Zeiss) with an oil-immersion 100× objective mounted on a piezo-controlled translator (Physik Instrumente). The magnetic field is generated by two coaxial coils (SBEA) with mu metal core (length, 40 mm; diameter, 26 to 88 mm; 750 spires). The coils are powered by a bipolar operational power supply amplifier 6A/36V (Kepco) controlled by a data acquisition module (National Instruments). The maximum field generated is 54 mT with a gradient less than 0.1 mT/mm over the sample. The self-organization of beads is first triggered with a constant field of 5 mT. Time-lapse images are recorded with an Orca Flash4 camera (Hamamatsu Photonics). The setup was heated at 37° C, without CO_2_ supplementation and is controlled by a custom LabVIEW interface that ensures the synchronicity between piezo position, magnetic field imposition, and image acquisition.

Cells with their cortex pinched between beads were imaged. For each film, the nominal magnetic field exerted by the coils was 5 mT (108 pN) and every 10 s the field was lowered to 1 mT (25 pN), then increased to 54 mT (1090 pN) in 1.5 s, then brought back to 5 mT (the values between brackets are the typical corresponding force values). This series of compression-relaxation was repeated 6 to 10 times per cell. In addition to the images, the precise time and magnetic field corresponding to each image were saved. Using Fiji’s plugin “Analyze Particle”, the center of all beads on each images of time-lapse was detected with a subpixel resolution^21^. Then using a home-made tracking algorithm written in Python (https://github.com/jvermeil-biophys/CortExplore_PublicVersion) the trajectories in 3D of the centers of the two beads pinching the cortex were detected, and the thickness of the pinched cortex was computed. The pinching force was also computed knowing the distance between the beads, the external magnetic field and the magnetization function of the beads. To characterize the cortex thickness, the median of the cortical thickness at 5 mT (nominal field) was measured; it correspond to a typical force of 108±29 pN. The elastic modulus at low stress was computed from the strain-stress curve corresponding to each compression using a model developed by Chadwick for this specific geometry^72^. Next, the slope of these strain-stress curves between 100 and 300 Pa was fitted.

### Immunofluorescence staining and quantification

Cells were seeded on micropatterned coverslips as described above and stained with MemBrite^®^ Fix before fixation according to the manufactures’ protocol. In brief, cells were washed twice with warm growth medium (37 ̊ C) and incubated for 10-15 min at 37 ̊ C and 5% CO_2_ before fixation. Next, cells were washed twice with PBS, incubated with the Prestaining solution (1:1000 in PBS) for 5 min at 37 ̊ C and stained with the Staining solution (1:1000 in PBS) for 5 min at 37 ̊ C.

Next, cells were fixed and permeabilized simultaneously by adding prewarmed (37 ̊ C) fixation buffer (2x) to growth medium and incubated for 20 min at RT. Fixation buffer (1x) contains 4% PFA (Thermo Fisher Scientific), 0.5% Tergitol (Serva), 20mM sucrose (Merck) in cytoskeleton buffer (10 mM MES, 140 mM KCl, 3 mM MgCl2, 2 mM EGTA, pH = 7.5). Cells were washed trice for 10 min with 0.05% Tergitol in TBS (Tris-Buffered Saline) and blocked for 1 hour with 2% BSA (Merck) in TBS. Primary antibody staining was performed over night at 4 ̊ C in blocking solution (anti-Phospho-MLC2 (Ser19), #3671, Cell signaling technology, 1:100). Cells were washed trice for 10 min with 0.05% Tergitol in TBS and incubated in blocking solution for 10 min. Secondary antibody staining was performed for 1 hour at RT in blocking solution (AF647 anti-rabbit #A21244, Thermo Fisher Scientific, 1:1.000; AF488 Phalloidin #A12379, Thermo Fisher Scientific, 1:1.000). Cells were washed trice for 10 min before mounting the coverslips on Marienfeld Superior Microscope slides with molds (Thermo Fisher Scientific) and imaged in TBS.

Cells were then imaged using an inverted confocal laser scanning microscope (Leica Stellaris 8^®^, HC PL APO CS2 40x / NA 1.10 water objective lens, white light laser) with LAS X software. For each cell one plane was recorded with a 4 μm set-off from the coverslip. Fluorescence intensities were analyzed using a custom-made Fiji macro, written with the kind help of Christian Tischer (EMBL Heidelberg). In brief, the cell was segmented based on the membrane staining, the mean fluorescence signal from F-actin, p-myosin and iMC-linker was measured in the cortical area (5, 10 and 15 pixels in width from the outer rim of the cell. See **Extended Data Fig. 1K**). Mean fluorescence intensities were normalized to the control for each replicate.

### mDia1 western blotting

For immune blotting, cells were lysed in RIPA Lysis and Extraction buffer (Thermo Fisher Scientific) supplemented with protease inhibitors (Roche) according to manufactures’ protocol. After removal of cell growth medium, cells were washed twice with cold PBS (4° C) before incubating in lysis buffer for 5 min on ice. Lysed cells were collected using a cell scraper. Cells were sonicated for 2 min to enhance lysis and centrifuged for 15 min at 14 Kg at 4° C. The supernatant was collected and treated with Bezonase (1:100) and 0.5 M MgCl2 (1:100) for 10 min at RT. Further, samples were denatured by mixing them with 4x Laemmli buffer (BioRad) containing 10% β-mercaptoethanol (Merck) and incubated for 5 min at 95° C. Proteins were separated in a SDS page using Nupage 4-18% Bis-Tris gels in running buffer.

Proteins were transferred by wet blotting (transfer buffer) on a methanol activated PDVF membrane. Successful protein transfer was verified by Ponceau red staining. For immunodetection, all washing steps were performed in TBST. The blot was blocked in 5% BSA in TBST and subsequently incubated with primary antibody (anti-mDia1, #20624-1-AP, Thermo Fischer Scientific, 1:1000, overnight at 4° C; GAPDH, #NB300-221, Novus Biologicals, 1:40.000, 1-2 hrs at RT). Secondary antibody staining was performed for 1 hour at RT in blocking solution (Donkey-Anti-Rabbit-HRP, #711-035-152, Jackson Immuno Research, 1:10.000; Goat-Anti-Mouse-HRP, #115-035-062, Jackson ImmunoResearch, 1:10.000). Proteins were detected using ECL western blotting substrate and a reader.

Western blots were quantified using gel analyzer plugin, available in Fiji. In brief, detected bands of interest were selected by the square tool and contained gray values are displayed as a histogram. To quantify the amount of protein, the area under the curve is integrated. For quantification, the signal of mDia1 was normalized to the GAPDH control and the mean of technical replicates was calculated and samples were normalized to untreated wild type (WT) samples.

### Cryo electron tomography

Micropatterns for cryo-electron tomography were designed as two 15 μm diameter circles with a separation of 10 μm in a single grid square. Micropatterns covered an area of 8 × 8 squares around the center of the cryo-TEM grids (Quantifoil R1/4 or R1/20, Au 200 mesh, SiO_2_ film) as previously described^31^. Briefly, grids were plasma treated on both sides for 60 s. Next, grids were incubated on droplets of poly-L-lysine grafted with polyethylene glycol (PLLg-PEG, Susos AG) at a concentration of 0.5 mg/ml in 10 mM HEPES pH 7.4, for 1 hour at RT on parafilm in a humid chamber. Following passivation, the grids were blotted with filter paper from the back, allowed to dry for few seconds and immediately placed on a drop of 4-benzoylbenzyl-trimethylammonium chloride (PLPP, Alvéole). Grids were then photo-patterned using the Primo module with a 375 nm laser (4.5 mW) as described above. Grids were promptly retrieved from the PLPP solution, washed first in milliQ water and then in PBS. For functionalization of the micropatterns, grids were incubated for 30 min at RT with fibronectin (50 μg/ml, Thermo Fischer Scientific) in PBS. Grids were then stored in PBS at 4° C in a humid chamber until use.

NIH 3T3 fibroblasts, at a concentration of 10^6^ cells/ml were seeded, allowed to settle and adhere for 20-30 min before non-attached cells were washed out. A Leica EM GP^®^ was utilized to vitrify cells 2 hrs after seeding. In the chamber, 1.5 μl of growth medium was directly applied to the backside of the grids. Blotting from the backside was performed for 2 s at 37° C, at 70% humidity. Grids were plunge-frozen in liquid ethane cooled by liquid nitrogen and stored until further usage.

For <u>cryo-focused ion beam (cryo-FIB) milling</u>, grids were clipped into an autogrid with a cut-out enabling milling at shallow angles. Mounted on a 45° pre-tilt shuttle, grids were transferred into an Aquilos Dual beam microscope (Thermo Fisher Scientific) via a load-lock system and maintained at −183° C throughout the next steps. Prior to cryo-FIB milling, a layer of organometallic platinum was applied by opening the gas injection system for 9 s at a stage Z position of 3-4 mm below the coincidence point. Subsequently, grids were sputter-coated with platinum for 20 s (1 kV, 10 mA, 10 Pa). In independent sessions, grids with 3-5 lamellae each were prepared at a milling angle of 15-18°. Grid squares with two cells attached in the desired geometry were micromachined in three steps of rough milling to a thickness of 5 μm at 1 nA ion beam current, 3 μm at 0.5 nA and 1 μm at 0.1 nA. Fine milling was performed at 100 and 50 pA to a target thickness of 200 nm. To render the lamellae conductive for TEM imaging, they were sputtered with platinum for 5 s (1 kV, 10 mA, 10 Pa) and transferred into sealed boxes in liquid nitrogen.

For <u>cryo-electron tomography</u>, autogrids containing lamellae were loaded into a Titan Krios^®^ microscope (Thermo Fisher Scientific) such that the axis of the pre-tilt introduced by cryo-FIB milling was aligned perpendicular to the tilt axis of the microscope. Tomograms were acquired at 300 kV on a K2 Summit direct detection camera (Gatan) operating in dose fractionation mode and utilizing a Quantum post-column energy filter operated at zero-loss (Gatan). Data were recorded using automated acquisition procedures in the SerialEM software^73^. Magnification of 42,000 (EFTEM) with a calibrated pixel size of 3.37 Å was used for data collection. Starting from the lamella pre-tilt, images were acquired at 2.0-4.0 μm under-focus, in 2° increments using a dose-symmetric tilt scheme^74^. A maximum of 125 e^−^/Å^2^ was used with a constant electron dose per tilt image. Tilt-series were collected with Volta potential phase plate (VPP, Thermo Fisher Scientific) with prior conditioning for 6 min. Cryo-electron tomography data analyzed in this work are summarized in **Supplementary Table 2**.

For <u>tomogram reconstruction</u>, tilt movie frames were aligned using a SerialEM plugin or in Warp (version 1.0.7)^75^. Tilt series alignment was performed in the IMOD (version 4.9) software package^76^, using fiducials (platinum particle contaminants on the lamellae surface) or patch-tracking. Aligned images were binned to the final pixel of 13.48 Å. Tomographic reconstruction was performed in IMOD.

Tomograms from **Fig. 2, Extended Data Fig. 3** and the **Supplementary Videos 1,3** were treated with an amplitude spectrum matching filter^77^.

<u>Actin segmentation</u> was performed with the automated filament tracing module in Amira on IMOD reconstructed tomograms, or tomograms treated with a spectrum matching, or gaussian filtered using a generic missing wedge-modulated cylinder with a radius of 4 nm and 40 nm length as a template. The traced filaments were manually curated in Amira, exported as a binary mask and the coordinates were resampled to equidistant points corresponding to the tomogram resolution (double the pixel size) using the Bspline fit in a python script.

<u>Membrane segmentation</u> was performed using the following workflow. An initial segmentation was generated using the TomoSegMemTV software package^78^. Parameters were tuned individually for each tomogram to achieve maximum membrane coverage for the plasma membranes which was determined by visual inspection. Voxels were then clustered using the connected component algorithm provided by the TomoSegMemTV package. The resulting clusters were processed using custom Python scripts. Each cluster was converted to a 3D point cloud (XYZ format) and imported into the Open3D library^79^ for further processing. Statistical outlier removal as implemented in Open3D was used to remove excess noise from the segmentations. In some cases, false positives (often microtubules or other intracellular membranes) were removed manually after visual inspection using Amira (Thermo Fischer). Subsequently, the DBSCAN algorithm was used to re-cluster and separate individual membrane sections from plasma membrane segments, which in some instances were classified as a single cluster (the analyzed tomograms always contained two plasma membrane segments form adjacent cells). Clusters corresponding to parts of the same membrane were then merged. A radial basis function fit as implemented in SciPy was then applied to point clouds of each plasma membrane to generate smooth, hole-free membrane surfaces for subsequent measurements.

<u>Quantitative analysis</u> of the cortex architecture was performed in MATLAB using the coordinates of the equidistant sampled actin points and the membrane segmentation binary mask as input. Each actin point in the tomogram was assigned to one of the two cells based on its distance to the segmented plasma membranes. Subsequently, the algorithms described by Jasnin *et al*.,^80^ were used to analyze the local-neighborhood of each actin point in each cell. Based on the peak in the 2D histogram of the nearest neighbor distance and relative orientation (**Extended Data Fig. 3b**), we identify actin bundles as two single actin points closer than 15 nm in distance with a relative orientation of less than 20°. Branch site candidates were initially identified as locations at the end of a filament fragment within a distance of 25 nm and angles between 50° and 90°, considering the 70° angle reported in literature ^81,82^. These candidates were further filtered, by determining the approximate point of intersection of the two filaments by extrapolating two line segments and only including those that pass each other within a maximal distance of 3 nm. Additionally, all candidates where the closest points of two filaments were end points were removed because visual inspection showed those instances being typically false positives. A subset of detected branches was visually inspected in the tomograms by 3 experts giving a 16.6% false positive rate. To ensure that the analysis was not affected by chosen parameters for inter-filament distance and angle, we varied these parameters and found the ratios of +iMC to -iMC were unaffected (**Extended Data Fig. 3c,d**).

To determine the membrane-to-cortex distance, for each resampled actin point, the distance to the closest membrane point was determined. The obtained distances were binned into 20 nm bins for each cell. The counts in each bin were normalized against the plasma membrane area of the cell, and the normalized counts were averaged over all cells to produce a final histogram (**Fig. 2b,c**). To calculate the angle between filaments and the membrane, the tangent plane of the membrane was defined as the plane orthogonal to the normal vector and determined using a script in python. For each actin point and its closest membrane point, the angle between the local orientation of the actin point and the tangent plane of the membrane point was calculated. The obtained angles were binned, normalized against the membrane surface area, and the normalized count was averaged across all tomograms per bin (**Extended Data Fig. 2f**).

Figures and movies were generated using IMOD, Chimera^83^ and Chimera X^84^.

### Statistical analysis

Statistical analyses were performed using R, while data visualization by both R and Adobe Illustrator^®^. Normality of data distribution was tested by Shapiro-Wilk test. Two-tailed t-test was used for normally distributed data. Otherwise, a non-parametric Wilcox test was used. In all box- and violin-plots, the lower and upper hinges correspond to the first and third quartiles (the 25th and 75th percentiles). The upper whisker extends from the hinge to the largest value, but no further than 1.5*IQR (distance between the first and third quartiles). The lower whisker extends from the hinge to the smallest value, but no lower than 1.5*IQR of the hinge. Data beyond the end of the whiskers are plotted as black dots. Black line corresponds to the median; each dot represents the mean of multiple measurements of single cells unless specified otherwise. In violin-plots the colored area reflects the probability distribution of the data points.

### Code availability

The Fiji macro for immunofluorescence quantification as well as the cryo-ET analysis code will be made available upon publication. Tomograms and segmentations generated in this work will be deposited to the EMDB and EMPIAR.

## Acknowledgements

We thank Jan Ellenberg, Stephan Grill and Anna Erzberger for critical reading of the manuscript. We thank Estela Sosa Osorio for manual curation of actin segmentation in cryo-electron tomograms, and Ievgeniia Zagoriy and Mukthi Ammai Sridharan Iyer for the visual inspection of actin branching points in cryo-electon tomograms. We thank the EMBL flow cytometry core facility, and the EMBL advanced light microscopy facility for support and advice, especially Christian Tischer for help with image analysis. We acknowledge the financial support of the European Molecular Biology Laboratory (EMBL) to J.M. and A.D-M., the Deutsche Forschungsgemeinschaft (DFG) grant DI 2205/2-1 and the Human Frontiers Science Program (HFSP) grant RGY0073/2018 to A.D-M., the Boehringer Ingelheim Fonds PhD fellowship and the Croucher Scholarship for Doctoral Study to D.C., and the EMBL interdisciplinary Postdoc (EIPOD) programme under Marie Curie Cofund Actions MSCA-COFUND-FP to M.S. and M.T-N.

## Author contributions

S.L., L.S., and A.D-M. conceived the project. S.L., L.S., J.M. and A.D-M. designed the experiments. M.T-N. performed the cryo-ET experiments under the supervision of J.M.. D.C., M.T-N. and M.S. analyzed the cryo-ET datasets under the supervision of J.M. and J.K.. J.V. performed and analyzed the magnetic pincher experiments under the supervision of M.P., O.D.R. and J.H.. C.J.C performed and analyzed the micropipette aspiration experiments. S.L. and L.S. performed and analyzed the rest of the experiments. S.L., L.S. and A.D-M. wrote the manuscript. All authors contributed to the interpretation of the data, and read and approved the final manuscript.

---

**Supplementary Table 1.** Transgenic cell lines used in this study.

| Cell line | Construct name | Description |
| --- | --- | --- |
| NIH 3T3 | CAezrin | constitutively active human ezrin (T567D)-mCherry |
| NIH 3T3 | iMC-linker | Lyn lipidation motif-mCherry-CH1/CH2 domain (human utrophin) |
| NIH 3T3 | iMC-6FP-linker | Lyn lipidation motif-mCherry-dark mCherry(x5)-CH1/CH2 (human utrophin) |
| NIH 3T3 | ABD | mCherry-CH1/CH2 domain (human utrophin) |
| mESCs | CAezrin | constitutively active human ezrin (T567D)-mCherry<sup>16</sup> |
| mESCs | iMC-linker | Lyn lipidation motif-mCherry-CH1/CH2 (human utrophin)<sup>16</sup> |
| mESCs | ABD | mCherry-CH1/CH2 domain (human utrophin)<sup>16</sup> |

**Supplementary Table 2.** Summary of tomography data.

| Condition | - iMC | + iMC |
| --- | --- | --- |
| Number of grids | 3 | 1 |
| Number of tomograms | 6 | 4 |
| Number of lamellae | 6 | 4 |
| Number of cells analyzed | 12 | 8 |
| Number of actin filaments | 1511 | 775 |

