## Supplementary materials for "The distance between the plasma membrane and the actomyosin cortex acts as a nanogate to control cell surface mechanics"

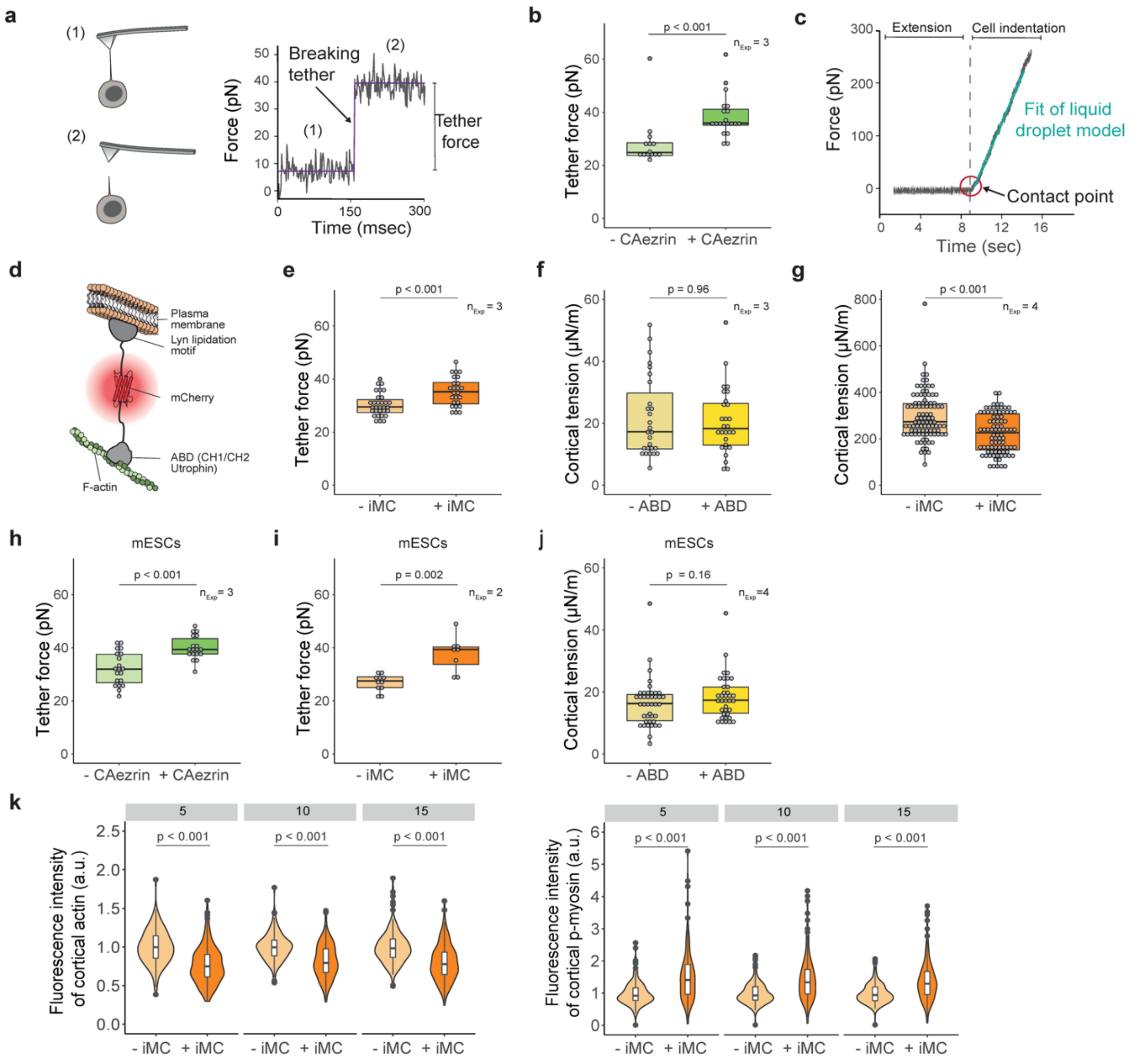


**Extended Data Fig. 1 | Biophysical characterization of the cell surface of NIH 3T3 fibroblasts and mESCs upon expression of different constructs. a**, Schematic representation of tether pulling with an atomic force spectrometer to quantify membrane-to-cortex tethering (left) and exemplary trace for tether pulling (right). Tether force is proportional to apparent membrane tension, and proxy for membrane-to-cortex tethering (see Methods). **b**, Membrane-to-cortex tethering upon expression of CAezrin in NIH 3T3 fibroblasts. **c**, Exemplary trace for nano-indentation. **d**, Schematic representation of iMC-linker. **e**, Membrane-to-cortex tethering upon expression of iMC-linker in NIH 3T3 fibroblasts. **f**, Cortical tension of NIH 3T3 fibroblasts expressing only the actin-binding domain of iMC-linker (ABD). **g**, Cortical tension measured with micropipette aspiration in NIH 3T3 fibroblasts in suspension upon expression of iMC-linker. **h,i**, Membrane-to-cortex tethering upon expression of CAezrin (h) or the iMC-linker (i) in mESCs. **j**, Cortical tension of mESCs expressing only the actin-binding domain of iMC-linker (ABD). **k**, Normalized mean fluorescence intensity of cortical F-actin (left) and p-myosin (right) with different segmentation depths (see methods). Each dot represents the mean of multiple measurements of a single cell. $n_{Exp}$ = number of independent experiments, a.u. = arbitrary units. Normality of data distribution was tested by Shapiro-Wilk test. Two-tailed t-test was used for normally distributed data. Otherwise, a non-parametric Wilcox test was used.

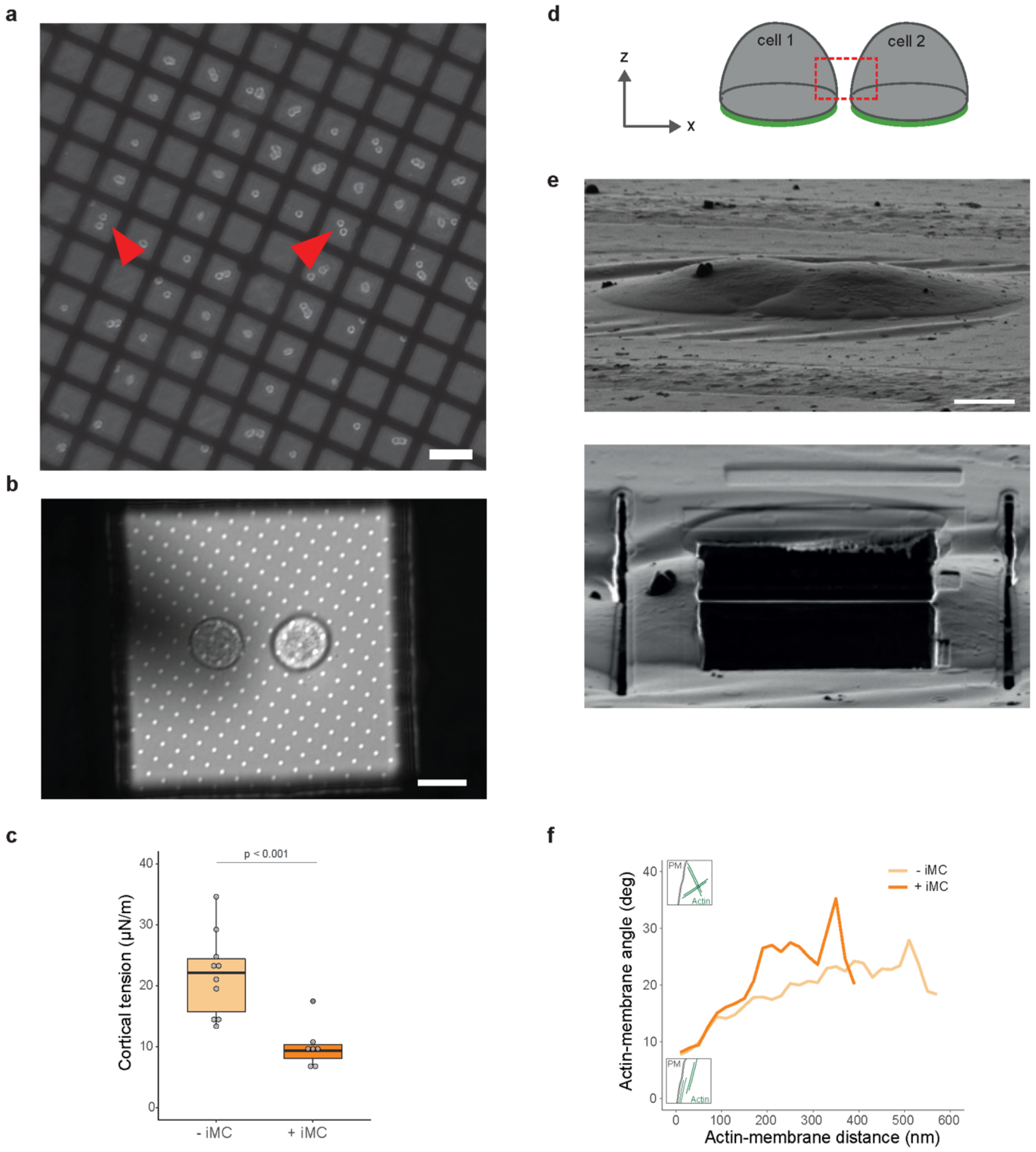


**Extended Data Fig. 2 | cryo-ET sample preparation. a**, Micropatterning on cryo-transmission electron microscopy (TEM) grids designed to accommodate two NIH 3T3 fibroblasts in close proximity to allow FIB-lamella generation at the cell surface of two neighboring cells. Red arrowheads indicate cell pairs in a suitable geometry for FIB milling. **b**, Bright field image of a representative cell pair seeded on a micropatterned grid square. **c,** Cortical tension measured by nano-indentation of cells on grids. Each dot represents the mean of multiple measurements of a single cell. Cells display a similar behaviour as on glass. See Fig. 1c for comparison. **d,** Lateral view schematic representation of a cell pair seeded on a grid square. Dashed red square marks area of FIB milling. **e**, Scanning electron microscopy images of a cell pair before (top) or after (bottom) FIB milling. Notably, cells underwent blotting before plunge-freezing, which leads to mild flattening. **f,** Average relative angle of actin filaments with respect to the plasma membrane measured from all tomograms in control or iMC-linker expressing NIH 3T3 fibroblasts. Inserts: schematics representing F-actin with large (top) and small (bottom) angle with respect to the plasma membrane. PM denotes the plasma membrane. Scale bars = 50μm in (a), = 10 μm in (b).

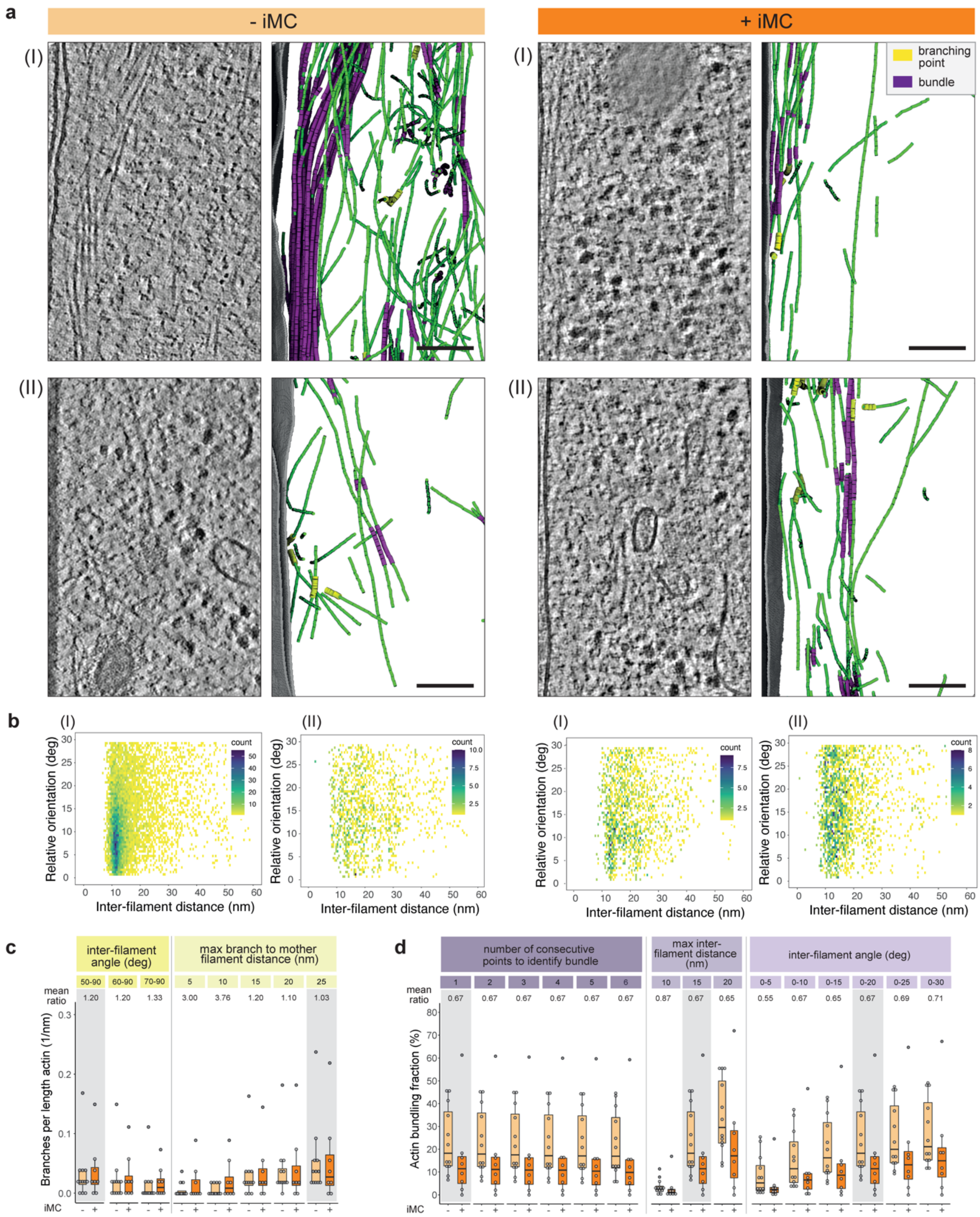


**Extended Data Fig. 3 | Supplemental cryo-ET data. a,** Two representative tomograms (I, II) and semi-automatic segmentation of actin filaments (green) and plasma membrane (grey) in control (- iMC) and iMC-linker expressing (+ iMC) NIH 3T3 fibroblasts, respectively. Segmented tomograms display branch sites (yellow segments) and actin in bundles (purple). Scale bars = 100 nm. **b,** Two-dimensional histogram of inter-filament distances and relative orientations for all points along the F-actin in tomograms shown in (a). A sharp peak at low angles and short distances indicates bundling. **c, d,** For the classification of actin branches (c) and actin bundles (d), parameters (angle and distance between filaments, and number of consecutive points for actin bundles) were varied to test the effect on the ratio between control (-iMC) to iMC-linker expressing (+iMC) NIH 3T3 fibroblasts. Actin branches (c) and the fraction of bundled actin (d) are normalized to the total length of actin per tomogram. Mean ratio = mean

(+iMC) divided by mean (-iMC); grey shadow marks the parameter combination used in the study (Fig. 2f,g).

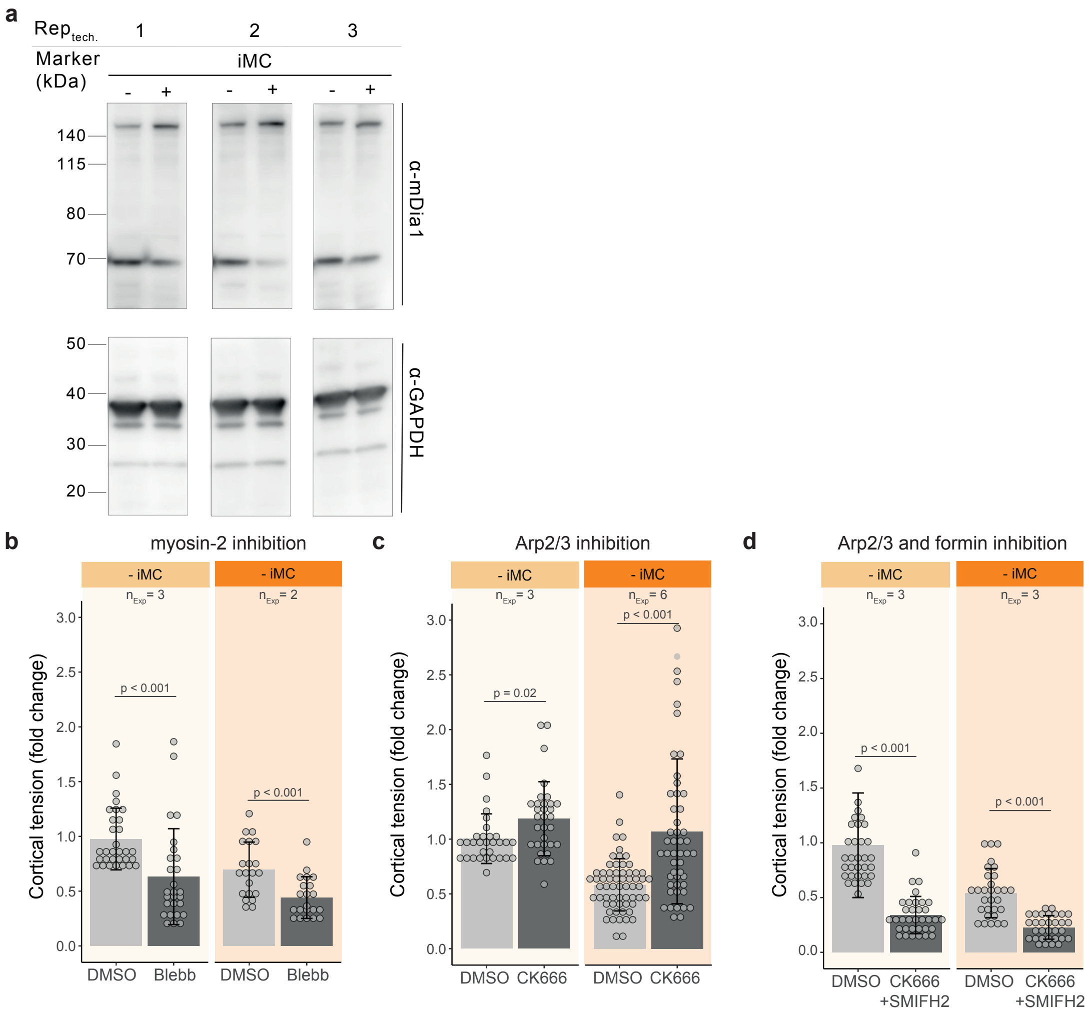


**Extended Data Fig. 4 | Extended drug perturbation panel to assess actomyosin regulators. a**, Representative uncut western blot for mDia1 and GAPDH. Shown are three technical replicates of one independent experiment. **b-d**, Relative cortical tension of NIH 3T3 fibroblasts upon treatment with the myosin inhibitor Blebbistatin (Blebb, b), treatment with the Arp2/3 complex inhibitor CK666 (c), or co-treatment with CK666 and the pan-formin inhibitor SMIFH2 (d). Tension values are normalized to the mean cortical tension of respective control cells (- iMC) treated with vehicle only (DMSO). Each dot represents the mean of multiple measurements of a single cell. $n_{Exp}$ = number of independent experiments. Normality of data distribution was tested by Shapiro-Wilk test. Two-tailed t-test was used for normally distributed data. Otherwise, a non-parametric Wilcox test was used.

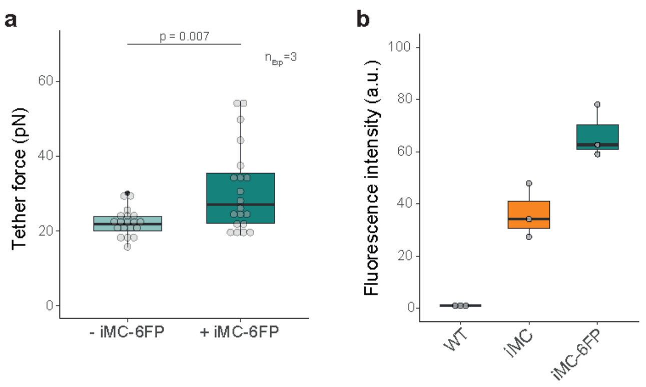


**Extended Data Fig. 5. | iMC-6FP-linker controls. a**, Membrane-to-cortex tethering upon expression of iMC-6FP-linker in NIH 3T3 fibroblasts. Each dot represents the mean of multiple measurements of a single cell. $n_{Exp}$ = number of independent experiments. **b**, Geometric mean of fluorescence intensities of iMC- or iMC-6FP-linkers. Each dot represents an independent experiment. WT: wild type NIH 3T3 fibroblasts, a.u. = arbitrary units. Normality of data distribution was tested by Shapiro-Wilk test. Two-tailed t-test was used for normally distributed data. Otherwise, a non-parametric Wilcox test was used.

**Supplementary Video 1.** Tomographic volume of the interface between two control NIH 3T3 fibroblasts. Tomogram corresponds to Figure 2a, and shown in the original orientation of the tomographic acquisition (tilt-axis aligned to the y-axis). Scale bar = 100 nm.

**Supplementary Video 2.** Corresponding 3D actin and plasma membrane segmentation of Supplementary Video 1 (green = F-actin, grey = membrane)**.** Detected actin bundles are highlighted in purple, actin branch sites are highlighted by yellow segments.

**Supplementary Video 3.** Tomographic volume of the interface between two NIH 3T3 fibroblasts upon expression of the iMC-linker. Tomogram corresponds to Figure 2a, and shown in the original orientation of the tomographic acquisition (tilt-axis aligned to the y-axis). Scale bar = 100 nm.

**Supplementary Video 4.** Corresponding 3D actin and plasma membrane segmentation of Supplementary Video 3 (green = F-actin, grey = membrane)**.** Detected actin bundles are highlighted in purple, actin branch sites are highlighted by yellow segments.
